# Engineered Prime Editors with PAM flexibility

**DOI:** 10.1101/2020.11.10.377580

**Authors:** Jiyeon Kweon, Jung-Ki Yoon, An-Hee Jang, Ha Rim Shin, Ji-Eun See, Gayoung Jang, Jong-Il Kim, Yongsub Kim

## Abstract

Although prime editors are a powerful tool for genome editing, which can generate various types of mutations such as nucleotide substitutions, insertions, and deletions in the genome without double-strand breaks or donor DNA, the conventional prime editors are still limited to its target scopes because of the PAM preference of the spCas9 protein. Here, we described the engineered prime editors to expand the range of their target sites using various PAM-flexible Cas9 variants.

---

Recently, Anzalone *et al.*^1^ developed prime editors (PEs), which can write new genetic information without double-strand breaks (DSBs) or donor DNA. PE2 utilizes an engineered *Moloney murine leukemia virus* (M-MLV) reverse transcriptase (RT) fused to a *Streptococcus pyogenes* Cas9 (spCas9)-H840A nickase and prime editing guide RNA (pegRNA) to manipulate the target locus. PEs allow the introduction of various types of mutations such as transition, transversion, insertion, and deletion in the various organisms^2, 3^; however, the NGG protospacer adjacent motif (PAM) preference of spCas9 restricts the targetable locus of PEs in the genome.

To overcome the limitation, we constructed PE2 variants having a preference for non-canonical PAMs using structure-guided engineered spCas9 variants, including the VQR and VRQR variants for NGA PAM, and the VRER variant for NGCG PAM from Joung’s group^4, 5^, the Cas9-NG variant for NG PAM from Nureki’s group^6^, and the SpG variant for NG PAM and SpRY variant for unconstrained PAM preferences from Kleinstiver’s group^7^. H840A mutations were further induced in each variant to generate nickases, and PE2 variants were constructed by changing the wild-type spCas9-H840A nickase to these nickases. We designated these variants as PE2-VQR, PE2-VRQR, PE2-VRER, PE2-NG, PE2-SpG, and PE2-SpRY.

To compare the activity of PE2 variants with that of wild-type PE2, we designed pegRNAs, which can target 35 genomic loci bearing NGN PAM. The pegRNAs had recommended length of primer binding site (PBS) and RT template (13 nt PBS and 11~14 nt RT templates) and were randomly designed to install insertions, deletions, and substitutions in the target genomic locus. We constructed the pegRNAs more efficiently using the single-strand DNA assembly method ^8^ (Supplementary Note 1). Using these pegRNAs, we examined the prime editing activity of six PE2 variants and wild-type PE2 in HEK293T cells (**Fig. 1a-c** and **Supplementary Table 1**). As expected, wild-type PE2 induced targeted mutations predominantly at the NGG PAM sites (active in 7 of 9 sites, up to 23.8% at the UBE3A-3+5G>C site) with some recognition of the NGA PAM sites (active in 4 of 9 sites, up to 22.9% at the VEGFA-4+1G>C site) and did not induce mutations at the NGC sites (active in 0 of 8 sites) or NGT sites (active in 0 of 9 sites). In contrast, the PE2-NG, PE2-SpG, and PE2-SpRY variants could edit NGN sites (PE2-NG: active in 24 of 34 sites, PE2-SpG: active in 26 of 35 sites, PE2-SpRY: active in 24 of 35 sites). In agreement with previous studies^6, 7^, the PE2-NG variant exhibited relatively lower activity at NGC PAM sites compared with NGD (where D is A, G, or T) PAM sites, and compared with other PE2 variants, the PE2-SpG variant showed the highest activity at NGH (where H is A, C, or T) PAM sites among the other PE2 variants (**Fig. 1c** and **Supplementary Fig.1**).

**Figure 1.**
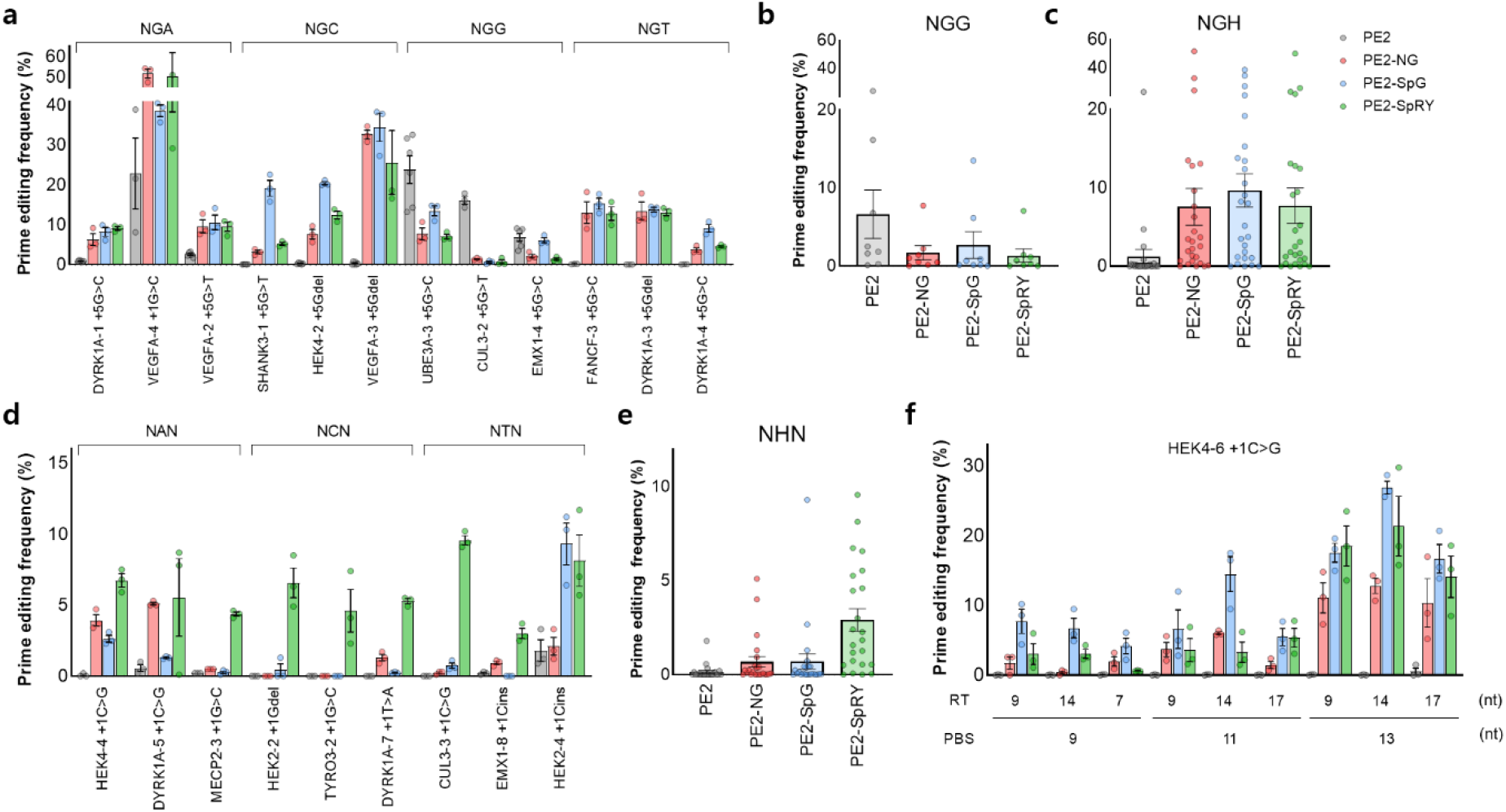
Prime editing in human cells by PE2 variants. **(a)** Representative prime editing activities across the NGA, NGC, NGG, and NGT PAM sites. Summary of prime editing activity at NGG PAM site, **(b)** and at NGH PAM site, **(c)**. **(d)** Representative prime editing activities at non-NGN PAM site (NAN, NCN, and NTN PAM). Summary of prime editing activity at NHN PAM site, **(e)**. **(f)** Prime editing activities at the HEK4-6 +1C>G site with pegRNAs of various lengths of PBS and RT templates. Mean ± s.e.m. of *n* = 3 independent biological replicates.

The PE2-VQR and PE2-VRQR variants exhibited higher activity at NGA PAM sites (both active in 5 of 7 sites, PE2-VQR: up to 32.7% at the VEGFA-2+5G>T site, PE2-VRQR: up to 30.2% at the VEGFA-1+5G>C site), and the PE2-VRER variant showed a preference for NGCG PAM compared with NGG PAM (up to 30.9% at the VEGFA-3+5Gdel site) (**Supplementary Fig. 2, 3**).

As the spCas9-SpRY variant can edit the genome in a near-PAMless manner, we investigated whether the PAM specificity of the PE2-SpRY variant was also relaxed. We additionally designed 23 pegRNAs bearing NHN PAM. The pegRNAs were randomly designed to install insertions, deletions, and substitutions in the target locus. The PE2-SpRY variant was active in 18 of 23 NHN PAM sites and induced prime editing up to 9.6% at the CUL3-3 +1C>G site, and other PE2 variants showed lower activity (PE2: active in 2 of 23 sites, PE2-NG: active in 5 of 23 sites, PE2-SpG: active in 4 of 23 sites) across NHN PAM sites (**Fig. 1d,e** and **Supplementary Table 1**). We examined a total of 58 pegRNAs targeting NNN PAM sites and found that the PE2-SpRY variant could manipulate genome in a PAM-independent manner (active in 43 of 58 sites). Collectively, these results showed that previously developed spCas9 variants could be successfully integrated into prime editing toolkits to relax the canonical PAM preferences.

Although we chose target sequences with high levels of indels generated by the spCas9 nuclease, prime editing activity varied from 0% to 51.7%, and many sites showed less than 0.5% activity (**Supplementary Table 1**). As previous studies described that the prime editing activity of wild-type PE2 is affected by the lengths of the PBS and RT templates, we examined whether the composition of pegRNA could affect the prime editing activity of PE2 variants. Two pegRNAs (HEK4-6+1C>G site with NGC PAM and VEGFA-4+1G>C site with NGA PAM) were selected and reconstituted to have various PBS and RT template lengths. At the HEK4-6+1C>G site, although wild-type PE2 was inactive with all pegRNAs, the PE2-NG, PE2-SpG, and PE2-SpRY variants were active and showed the highest prime editing activity (12.8%, 26.9%, and 21.4%, respectively) by pegRNAs with the 13 nt PBS and 14 nt RT templates (**Fig. 1f**). Longer PBS were preferred for all three PE2 variants (average of 3.29% with 9 nt PBS and 16.6% with 13 nt PBS); however, there was no preference in terms of the lengths of the RT templates. These preferences were also observed at the VEGFA-4+1G>C site (**Supplementary Fig. 4**). As prime editing activity varied from 0% to 51.7% depending on the lengths of the PBS and RT templates, optimization of pegRNAs is essential to maximize the activity of the PE2 variants.

Previously, compared with Cas9 nuclease, wild-type PE2 has lower off-target effects at already known Cas9 off-target sites^1^. As there have been no further advances in the technology to analyze PE2-dependent off-target effects, we chose several sites to validate the potential off-target effects of PE2 variants as follows: 1) HEK4+2G>T site and its off-target locus, which is known as an off-target site for wild-type PE2; 2) FANCF-4+5Cins, EMX1-4+5G>C, and MECP2-3+1G>C sites and their off-target locus, which are known as off-target editing sites for the SpCas9-SpRY variant; 3) CUL3-3 +1C>G and HEK2-4 +1Cins sites, which showed high activity with the PE2-SpRY variant at NHN PAM sites, and their homologous locus. Using targeted-deep sequencing, we found that the PEs including wild-type PE2, PE2-SpG, and PE2-SpRY induced much lower off-target editing compared with their nucleases (spCas9, spCas9-SpG, and spCas9-SpRY) at these sites (increased specificity ratio up to 36.8-fold) (**Supplementary Fig. 5**). The high specificities of the PEs may be attributed to the requirement of additional base pairing with the PBS and RT templates to edit the genome.

Anzalone et al.^1^ demonstrated an increase in prime editing activity by inducing additional nicks on non-edited strand, which was named PE3 or PE3b. To examine whether additional nicks affect the prime editing activity of PE2 variants, we constructed additional gRNAs to incorporate PE3, which can induce additional nicks distant from pegRNA-induced nicks, or PE3b, which can induce nicks on the non-edited strand only after the resolution of the edited strand flap. At the HEK2-2+1Gdel site, PE3 improved prime editing activity up to 2.8-fold (6.6% to 18.7%), and PE3b improved prime editing activity up to 2.2-fold (6.6% to 14.8%) (**Fig. 2a**). At the CUL3-3+1C>G site, PE3 increased prime editing up to 2.9-fold (12.0% to 34.4%), and at the HEK4-4+1G>C site, PE3b increased prime editing up to 2.1-fold (5.4% to 11.4%) (**Fig. 2b** **and Supplementary Fig. 6**). Interestingly, we could not detect any increase in indels at these three sites with the PE2-SpRY variant when using different doses of gRNAs with PE3 or PE3b. We believe that the nicking activity of spCas9-H840A variants may affect the indel frequencies of PE3 or PE3b systems. Overall, we found that PE3 and PE3b could successfully increase the prime editing activity of PE2 variants without excess indel formation.

**Figure 2.**
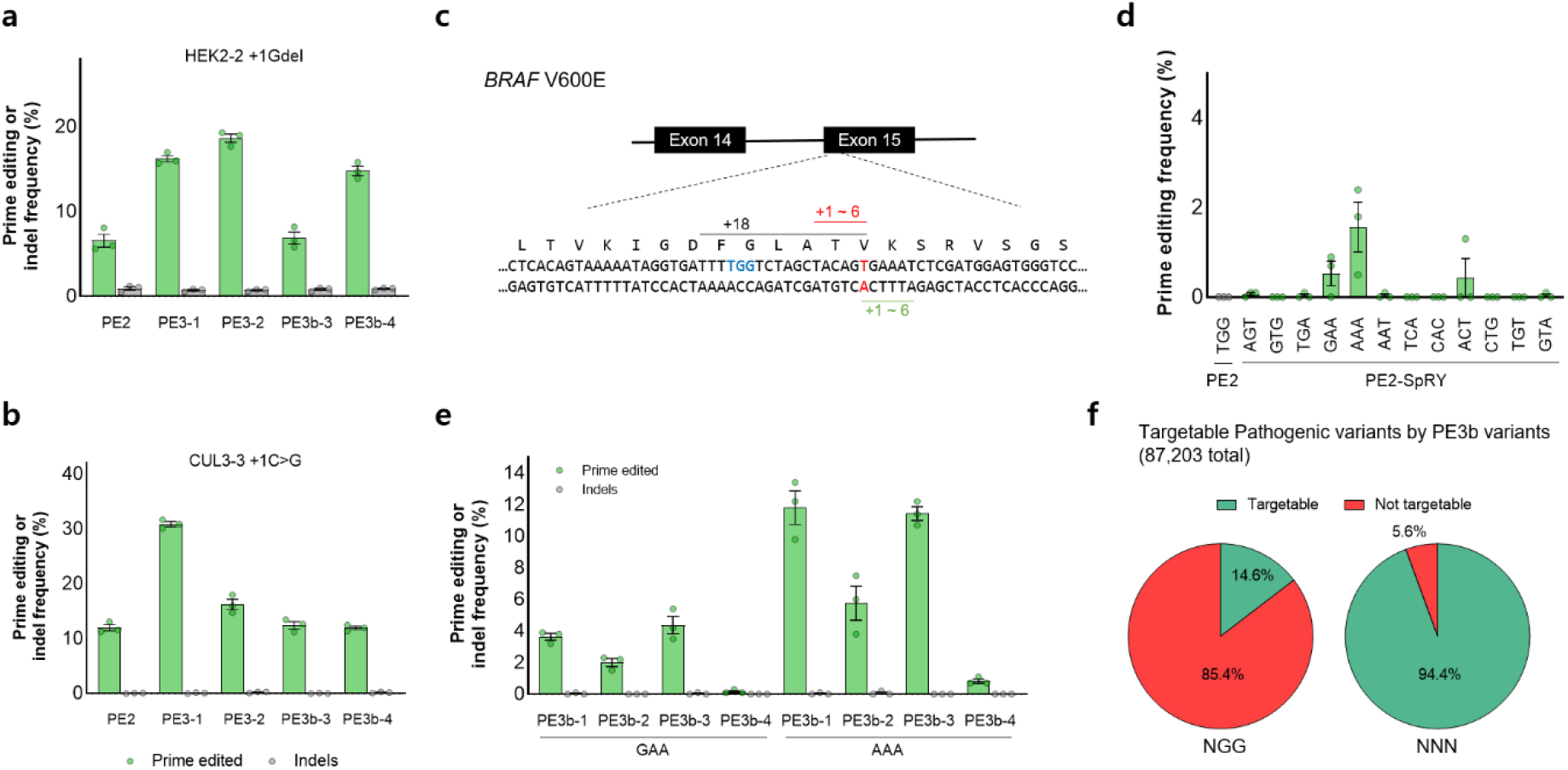
Prime editing by PE3 and PE3b systems, and the BRAF V600E mutation by the PE2-SpRY variant. Comparison of prime editing and indel frequencies at the HEK2-2 +1Gdel site, **(a)**, and at the CUL3-3 +1C>G site, **(b)**. **(c)** The schematic overview of prime editing for the BRAF V600E mutation. The target thymine nucleotide is highlighted in red, and the closest PAM sequence of wild-type PE2 is indicated in blue, which can induce nick 18 nt away from that target thymine nucleotide. The pegRNAs of PE2-SpRY were designed to induce nicks 1-6 nt in both directions from the target thymine. **(d)** Prime editing activities of PE2 and PE2-SpRY at the BRAF V600E site. **(e)** Prime editing activity of the PE2-SpRY by PE3 and PE3b systems at the BRAF V600E site. **(f)** Comparison of prime editable pathogenic variant using NGG or NNN PAM in the PE3b system.

BRAF V600E (which is caused by a T:A to A:T transversion mutation) is a well-known driver mutation in various cancers and diseases and a therapeutic target for anti-cancer drugs^9^ (**Fig. 2c**). Despite the broad ability of wild-type PE2 to introduce various pathogenic mutations, the distance between the pegRNA nick site and BRAF V600E target locus is 18 nt, which could not be efficiently targeted with wild-type PE2 (**Fig. 2d**). To compare the targeting efficiency of the wild-type PE2 and PE2-SpRY variant, we constructed pegRNAs with single-base resolution around the BRAF V600E target locus and examined their prime editing activity in HEK293T cells. We found that PE2-SpRY improved prime editing activity up to 6.7-fold compared with the activity of wild-type PE2 (0.27% to 1.80%); however, there was no significant increase in prime editing activity by the optimization of the lengths of PBS and RT templates (**Supplementary Fig. 7**). Then, we chose two active pegRNAs, BRAF-GAA and BRAF-AAA, and assessed the PE3 and PE3b systems; we found that PE3b significantly improved prime editing activity up to 11.8 % with BRAF-AAA pegRNA and BRAF-PE3b-1 gRNA (**Fig. 2e** **and Supplementary Fig. 8**).

In summary, we generated various types of PE2 variants with PAM flexibility and analyzed the prime editing activity of the variants at 71 target sites. The PE2-SpG variant may be suitable for targeting NGH PAM sites; specifically, the PE2-SpG variant was more active than the PE2-NG variant at NGC PAM sites. The PE2-SpRY variant induced prime editing without PAM restriction, albeit reduced in relative activity. The PE2-SpRY variant could offer increased prime editable sites per human pathogenic variant (covered 94.4% of pathogenic variants) through the PAM flexibility and could also increase the number of PE3b applicable sites (up to 28.8 sites per pathogenic variant) for improving prime editing efficiency without unwanted mutations (**Supplementary Fig. 9 and 10**). Recent studies are helpful for design of pegRNAs and analysis of PE-meditated off-target effects^10, 11^. Our constructs have been shared to Addgene for the research community.

## Data availability

All data supporting the findings of this study are available in the article or in Supplementary information files or are available from the corresponding author upon request. The deep sequencing data have been deposited in an NCBI BioProject database (PRJNA675696). All plasmids used in this study are available through Addgene.

## Acknowledgments

This work was supported by the National Research Foundation of Korea (grants 2020R1F1A1075508, 2017M3A9B4062419, and 2018R1A5A2020732 to Y.K.)

## Author contributions

Y.K. supervised the research. J.K., A-H.J., HR.S., and J-E.S. carried out experiments. J.K. and Y.K. performed data analysis. J-K.Y. analyzed ClinVar database. J-K.Y. and J-I.K provide conceptual advice. Y.K. and J.K. wrote the manuscript.

