## Supplementary Information for "Engineered Prime Editors with PAM flexibility"

**Table of contents**

Materials and Methods

Supplementary Note 1. Methods for pegRNA subcloning using ssDNA assembly.

Supplementary Figure 1. Prime editing activities of PE2 variants at NGC, NGA, and NGT PAM sites.

Supplementary Figure 2. Prime editing activities of PE2-VQR and -VRQR variants.

Supplementary Figure 3. Prime editing activities of the PE2-VRER variant.

Supplementary Figure 4. Prime editing activities at the VEGFA-4 +1G>C site with pegRNAs of various lengths of PBS and RT template.

Supplementary Figure 5. Off-target analysis of PE2 variants.

Supplementary Figure 6. Prime editing using PE3 and PE3b systems at three target sites.

Supplementary Figure 7. Prime editing activities at the BRAF V600E site with pegRNAs of various lengths of PBS and RT template.

Supplementary Figure 8. Prime editing using PE3 and PE3b systems at the BRAF V600E site.

Supplementary Figure 9. Fraction of human pathogenic variants which could be targetable with the PE2 or PE3b system.

Supplementary Figure 10. Numbers of target site per targetable variants in the PE2 and PE3b systems.

Supplementary Table 1. Summary of all prime editing activity analyzed in this study.

Supplementary Table 2. List of target sites used in this study.

Supplementary Table 3. List of off-target sites in this study.

Supplementary Table 4. Analysis of targetable pathogenic variants using PE variants.

### Materials and Methods

#### Plasmids DNA and pegRNAs

To construct PE2 variants, we used pCMV-PE2 (addgene #132775) plasmids DNA as vector DNA and the coding sequences of spCas9 variants were obtained by PCR amplification from NG-ABEMAX (NG variant, addgene plasmids # 124163), RTW3520 (VQR variant, addgene plasmids #139990), RTW3160 (VRER variant, addgene plasmids #139991), RTW3161 (VRQR variant, addgene plasmids #139992), RTW4177 (SpG variant, addgene plasmids #139998), and RTW4830 (SpRY variant, addgene plasmids #139989). All the PE2 variants used in this study were available from Addgene. The pegRNAs were cloned into the pU6-pegRNA-GG-acceptor (addgene plasmids #132777) and the spacer sequences and PBS plus RT template sequences were listed in Supplementary Table 2.

#### Mammalian cell culture and transfection

HEK293T cells (ATCC CRL-11268) were maintained Dulbecco's modified Eagle's medium (DMEM) supplemented with 10% fetal bovine serum (FBS) and 1% penicillin/streptomycin (Welgene). Cells were not tested for mycoplasma contamination. HEK293T cells were seeded onto TC-treated 96-well plate (Corning) at $3x10^4$ cells per well density one day before transfection. Transfection was conducted using 0.6 µl Lipofectamine 2000 (Thermo Fisher Scientific) according to the manufacturer's protocol at approximately 60% cell confluency. For PE2 experiments, 300 ng PE2 variants plasmids and 100 ng pegRNA plasmids DNA were used. For PE3 or PE3b experiments, 270 ng PE2 variants plasmids, 30 ng gRNA plasmids, and 100 ng pegRNA plasmids DNA were used for low dose experiments, and 200ng PE2 variants, 100 ng gRNA plasmids, and 100ng pegRNA plasmids DNA were used for high dose experiments. The transfected cells were incubated at 37°C for 3 days and genomic DNA was prepared by directed lysis the cells using the lysis buffer [10 mM Tris-HCl with pH 7.5, 0.05% SDS, 100 µg/ml proteinase K (Qiagen)]. The cell lysate was incubated 56°C for 30 min, followed by 99°C 15min incubation. In off-target analysis experiments, $1.5x10^5$ HEK293T cells were seeded onto TC-treated 24-well plate (Corning) and 1.5 µg of PE2 variants plasmids and 500 ng pegRNA plasmids DNA were transfected, and genomic DNA was isolated using the DNeasy Blood & Tissue kit (Qiagen) according to the manufacturer's protocol.

#### Targeted-deep sequencing and data analysis

The target sites were amplified by a total of three rounds of PCR and subjected to the Illumina Miniseq or iSeq 100 as previously described[1]. Briefly, 3 µl of cell lysate or 1 µl of isolated genomic DNA was subjected to the 1st round PCR and then, 1 µl of 1st PCR product was used in the 2nd round PCR. The Illumina TruSeq HT dual index adapter sequences were attached with index PCR primer pairs using 1 µl of 2nd round PCR product. The size of PCR amplicon was confirmed in 2% agarose gel and the amplicons were subjected to 150-bp paired-end sequencing using Illumina Miniseq or iSeq 100. The paired-end reads were joined using fastq-join tool (https://github.com/brwnj/fastq-join). Targeted-deep sequencing analysis was performed using MAUND[2] (https://github.com/ibs-cge/maund ) and all results are confirmed by Cas-Analyzer[3] (http://www.rgenome.net/cas-analyzer/). Substitutions and indel frequencies were quantified as the percentage of total sequencing reads.

#### Analysis of targetable pathogenic variants in ClinVar database

All possible pegRNAs for correcting pathogenic variants in ClinVar database (https://ftp.ncbi.nlm.nih.gov/pub/clinvar/) were designed by using the pipeline (https://gitlab.com/sanjanalab/primeediting) developed by Morris and colleagues[4] with modifications. Briefly, we downloaded all variants in ClinVar (released date: 2020/08/10) and selected the variants with "pathogenic" identifier. Only single base-pair substitutions and insertions and deletions of 10 base pairs or less were calculated as targeted variants. We designed and counted the number of all possible probes for PE2 and PE3b separately with either NGG, NG (PAM for PE2-NG and PE2-SpG, or NNN (PAM for PE2-SpRY) for each pathogenic variant. The length of PBS and RT template were fixed to 13 and 16 nucleotides, respectively. The secondary sgRNAs for PE3b were designed separately and matched with possible PE2 probes complementary to the opposite strand.

**Supplementary Note 1.** Methods for pegRNA subcloning using ssDNA assembly.

To improve the throughput of pegRNA construction, we used the ssDNA assembly method instead of the Golden Gate (GG) assembly method. In comparison with Golden Gate assembly, ssDNA assembly has no oligonucleotide annealing step in a thermocycler and does not require the phosphorylation of gRNA scaffold oligonucleotides. The ssDNA assembly method has a similar ratio of red/white colonies as shown below, where GG-RT is the incubation at room temperature for 10 min, and GG-Cycle is the cycle between 5 min at 16°C and 5 min at 37°C for 8 cycles, as described by Anzalone et al. We also determined the cloning efficiency by the targeted-deep sequencing of pooled transformants and found that ssDNA assembly has a slightly higher success rate in pegRNA cloning compared with conventional methods. The detailed protocol of pegRNA cloning is described below.

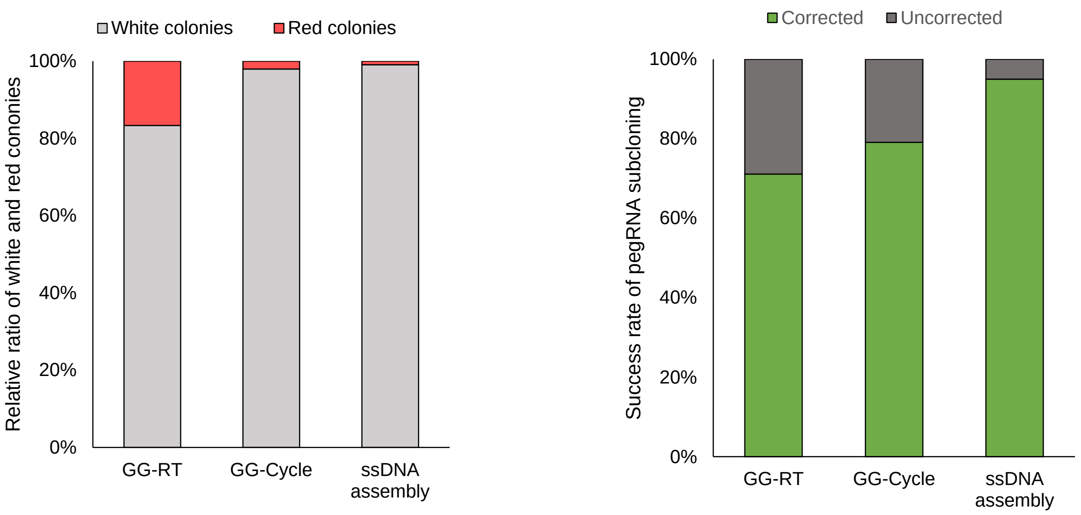


1. Oligonucleotide preparation for pegRNA cloning

    a. Oligonucleotides for spacer (target-specific component)
    The 18 nt 5' overhang (5'-tggaaaggacgaaacacc-3') and 3' overhang (5'-gttttagagctagaaata-3') are attached to the desired spacer sequence as follows:
    5'- tggaaaggacgaaacaccNNNNNNNNNNNNNNNNNNNNgttttagagctagaaata-3',
    where N×20 is the target spacer sequence without PAM, and the spacer sequence must begin with a guanine nucleotide. If the spacer sequence does not begin with a guanine nucleotide, an additional guanine nucleotide should be added for pegRNA transcription by the U6 promoter.

    b. Oligonucleotides for RT template & PBS (target-specific component)
    The 18 nt 5' overhang (5'-gtggcaccgagtcggtgc-3') and 3' overhang (5'-ttttttaagcttgggcc-3') are attached to the desired spacer sequence as follows:
    5'-gtggcaccgagtcggtgcNNNNNNNNNNNNNNNNNNNNttttttaagcttgggcc-3',
    where Ns are the desired RT template & PBS sequence.

    c. Oligonucleotides for pegRNA scaffold sequence (common component)
    The reverse complementary sequence of gRNA scaffold is as follows:
    5'-gcaccgactcggtgccacttttcaagttgataacggactagccttattttaacttgctatttctagctctaaaac-3'

2. Vector preparation for pegRNA cloning

    The pU6-pegRNA-GG-acceptor (addgene, plasmid #132777) plasmid DNA is digested with

BsaI-HFv2 (NEB R3733S) restriction enzyme as previously described.

3. pegRNA construction by ssDNA assembly
   Each component is mixed for ssDNA assembly as follows:

| | |
|---|---|
| Vector DNA | 50 ng |
| Oligonucleotides for spacer | 1 µl of 100 nM |
| Oligonucleotides for RT template & PBS | 1 µl of 100 nM |
| Oligonucleotides for gRNA scaffold | 1 µl of 100 nM |
| 2× HiFi DNA Assembly Master Mix (NEB #E2621S) | 5 µl |
| $H_2O$ | up to 10 µl |

   The reaction mixture is incubated at 50°C for 30 min, and an appropriate amount is transformed using competent cells.

The following scheme summarizes the ssDNA assembly for pegRNA subcloning protocol.

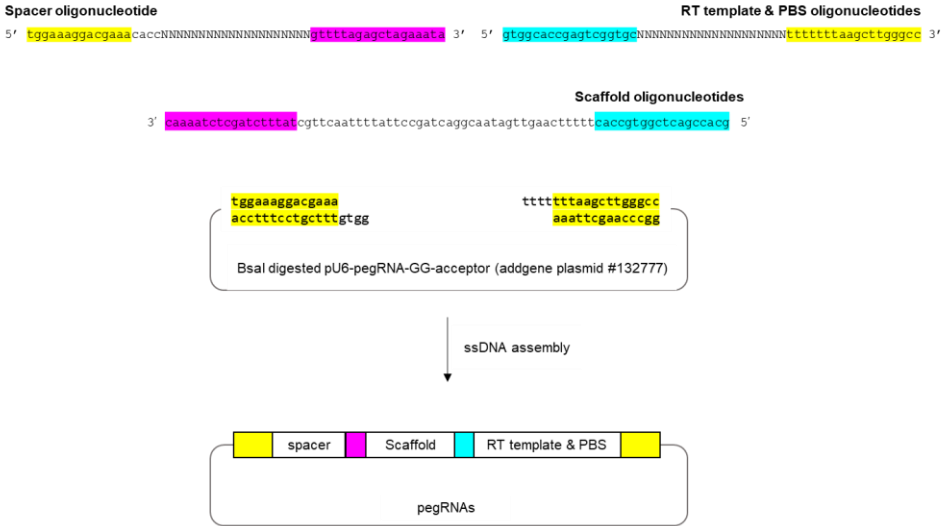

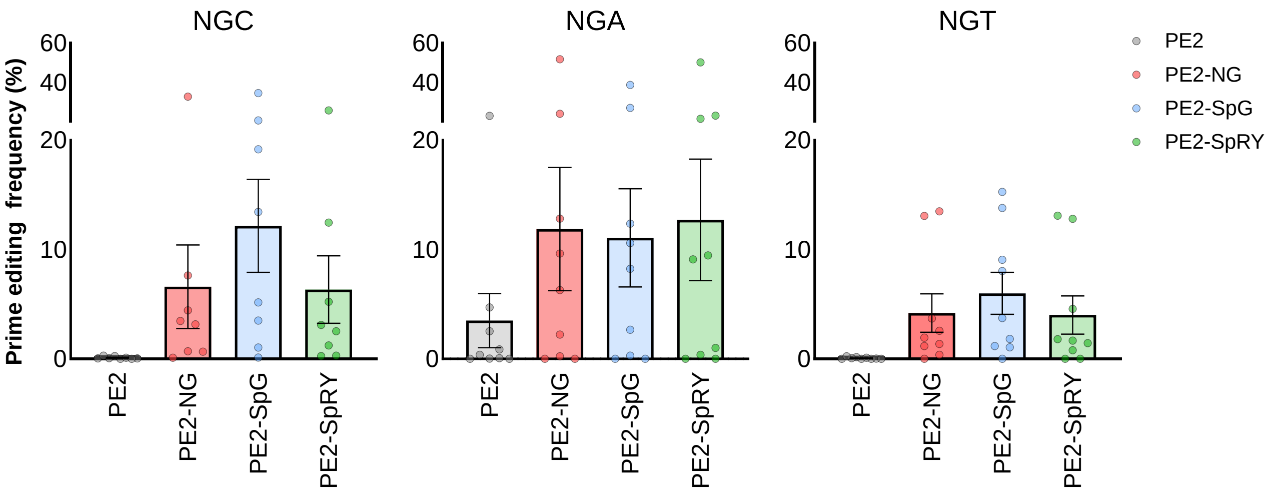


**Supplementary Figure 1.** Prime editing activities of PE2 variants at NGC, NGA, and NGT PAM sites. The PE2-SpG variant showed average 12.2% and 6.0% prime editing activity across the 8 sites with NGC PAM and the 9 sites with NGC PAM, respectively. The wild-type PE2 showed average 2.5% prime editing activities across 9 sites with NGA PAM. The numerical values of the prime editing activity are described in Supplementary Table 1. Mean ± s.e.m. of $n$ = 3 independent biological replicates.

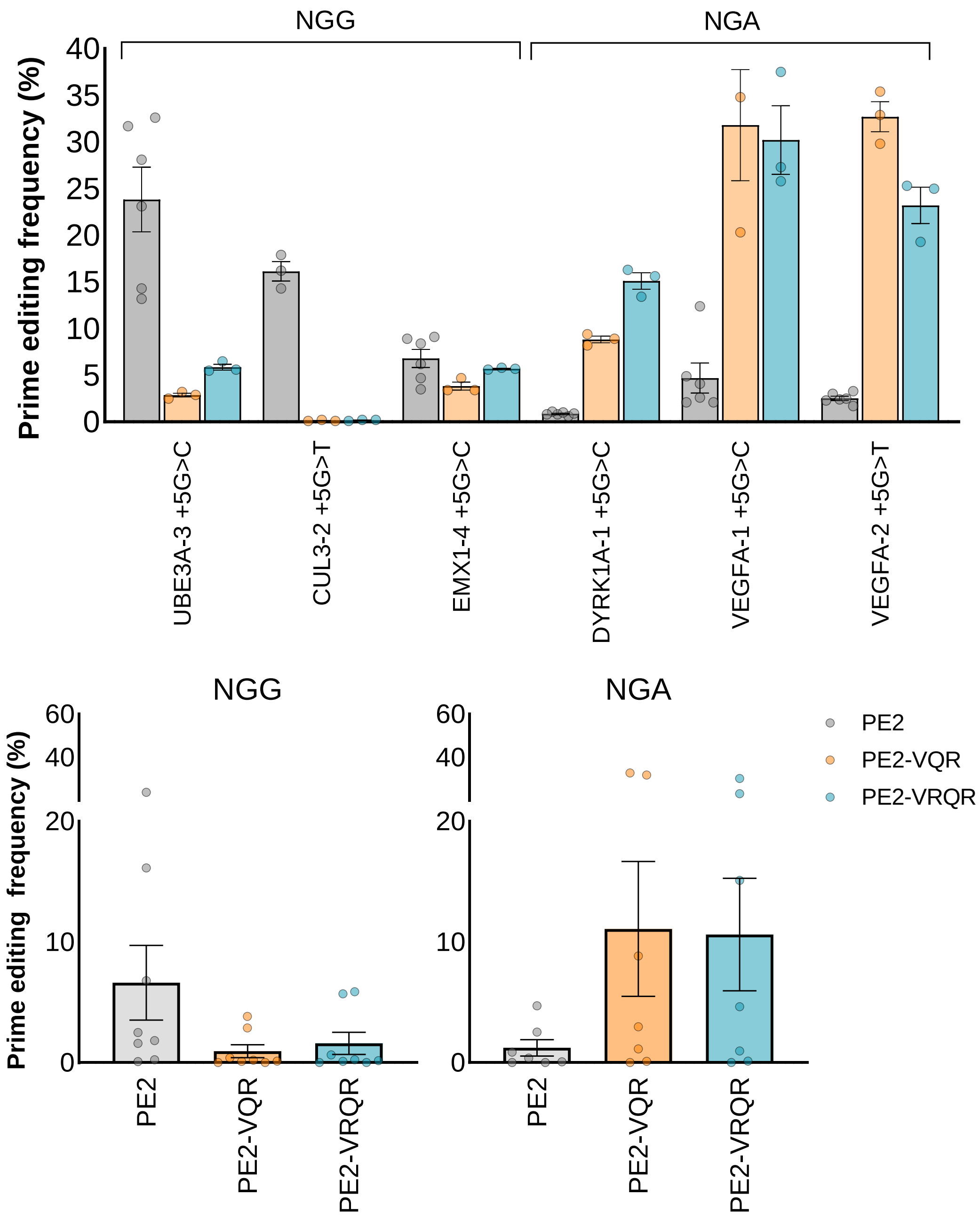


**Supplementary Figure 2.** Prime editing activities of PE2-VQR and -VRQR variants. The PE2-VQR and -VRQR variants showed average 11.1% and 10.7% prime editing activity across 7 sites with NGA PAM, respectively. The numerical values of the prime editing activity are described in Supplementary Table 1. Mean ± s.e.m. of $n = 3$ independent biological replicates.

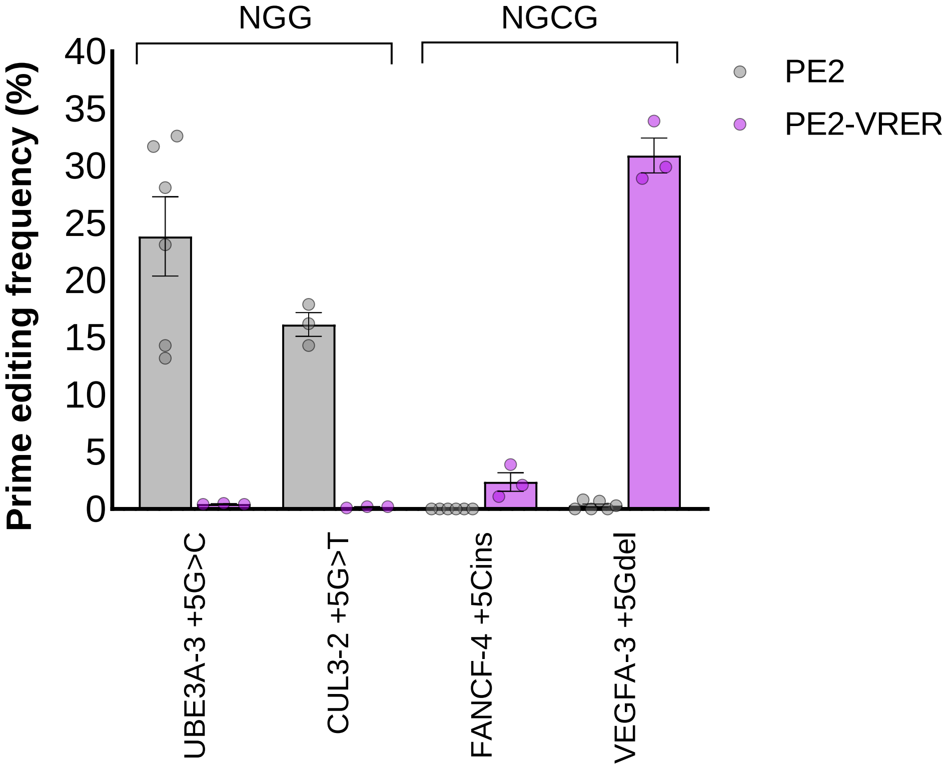


**Supplementary Figure 3.** Prime editing activities of the PE2-VRER variant. The PE2-VRER variant showed average 30.9% prime editing activity at VEGFA-3+5Gdel site with the NGCG PAM. The numerical values of the prime editing activity are described in Supplementary Table 1. Mean ± s.e.m. of $n$ = 3 independent biological replicates.

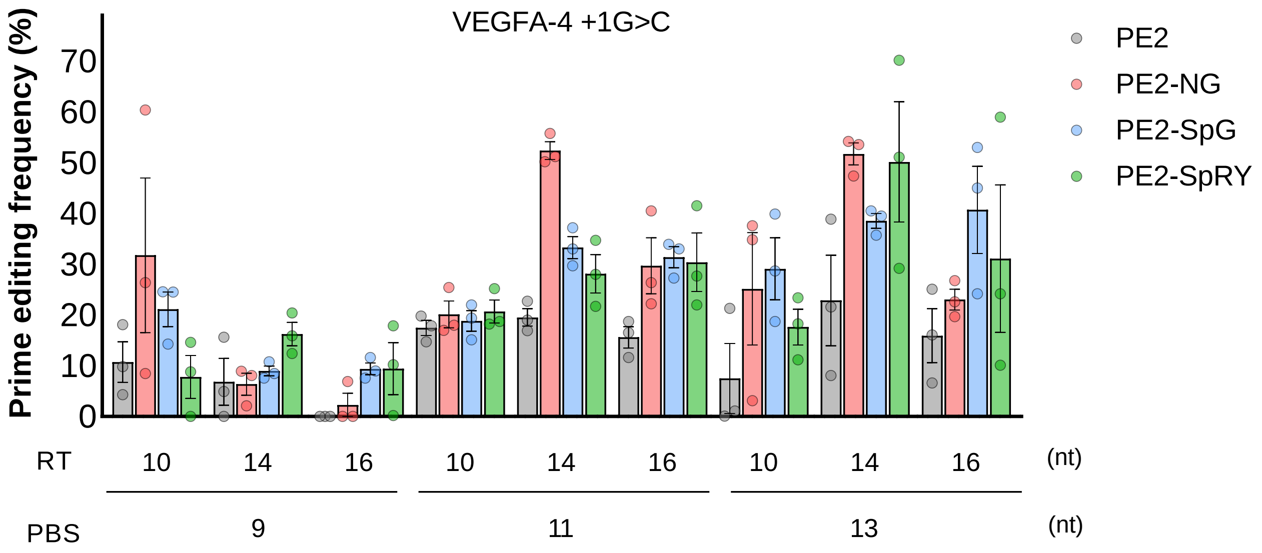


**Supplementary Figure 4.** Prime editing activities at the VEGFA-4 +1G>C site with pegRNAs of various lengths of PBS and RT template. The numerical values of the prime editing activity are described in Supplementary Table 1. Mean ± s.e.m. of $n = 3$ independent biological replicates.

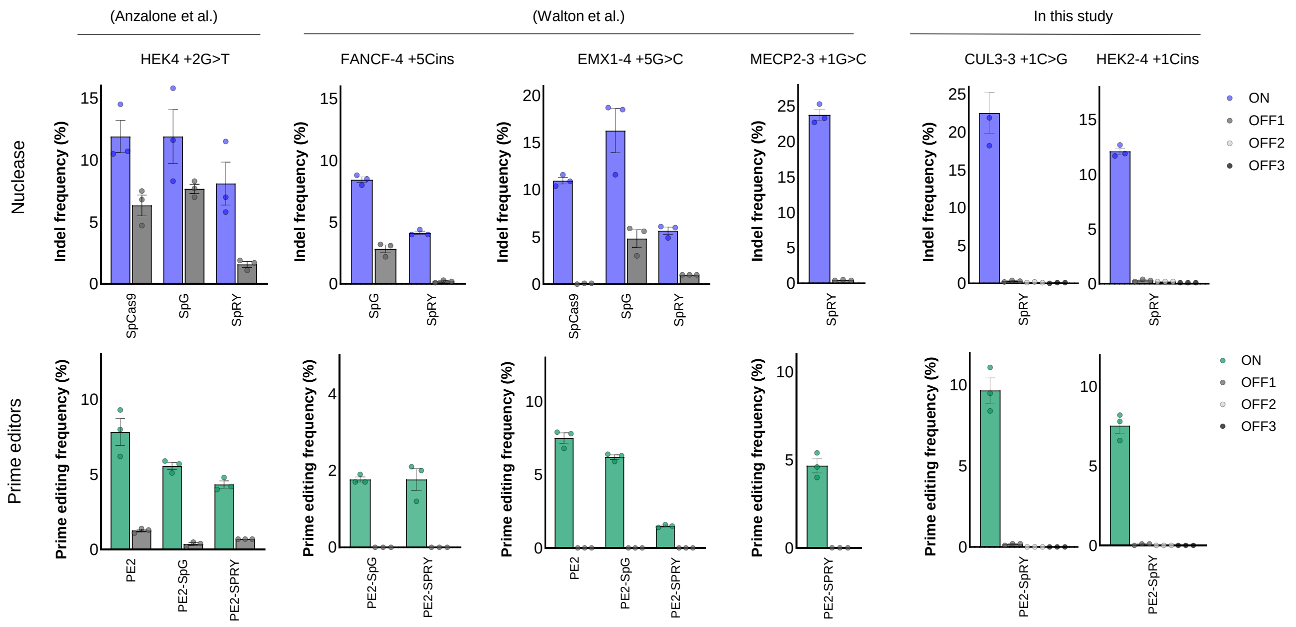


| Specificity ratio | HEK4 +2G>T | | | FANCF-4 +5Cins | | EMX1-4 +5G>C | | | MECP2-3 +1G>C | CUL3-3 +1C>G | | | HEK2-4 +1Cins | | |
|---|---|---|---|---|---|---|---|---|---|---|---|---|---|---|---|
| | | | | | | | | | | SpRY | | | SpRY | | |
| | WT | SpG | SpRY | SpG | SpRY | WT | SpG | SpRY | SpRY | OFF1 | OFF2 | OFF3 | OFF1 | OFF2 | OFF3 |
| Nulcease | 1.88 | 1.55 | 5.17 | 2.98 | 20.67 | 164.50 | 3.37 | 5.67 | 54.85 | 50.00 | 225.00 | 112.50 | 40.33 | 121.00 | 60.50 |
| PE2 | 6.18 | 6.19 | 15.18 | >35 | >35 | >150 | >124 | >30 | >93 | 57.60 | >192 | >192 | 113.00 | >150 | >150 |

**Supplementary Figure 5.** Off-target analysis of PE2 variants. We assessed six on-target sites and their ten off-target sites with nuclease (spCas9, spCas9-SpG, spCas9-SpRY) and PEs (wild-type PE2, PE2-SpG, and PE2-SpRY) listed in Supplementary Table 3. The specific ratio in the bottom means the relative mutation frequency of on-target sites compared to that of off-target sites. The numerical values of the prime editing activity are described in Supplementary Table 1. Mean ± s.e.m. of $n$ = 3 independent biological replicates.

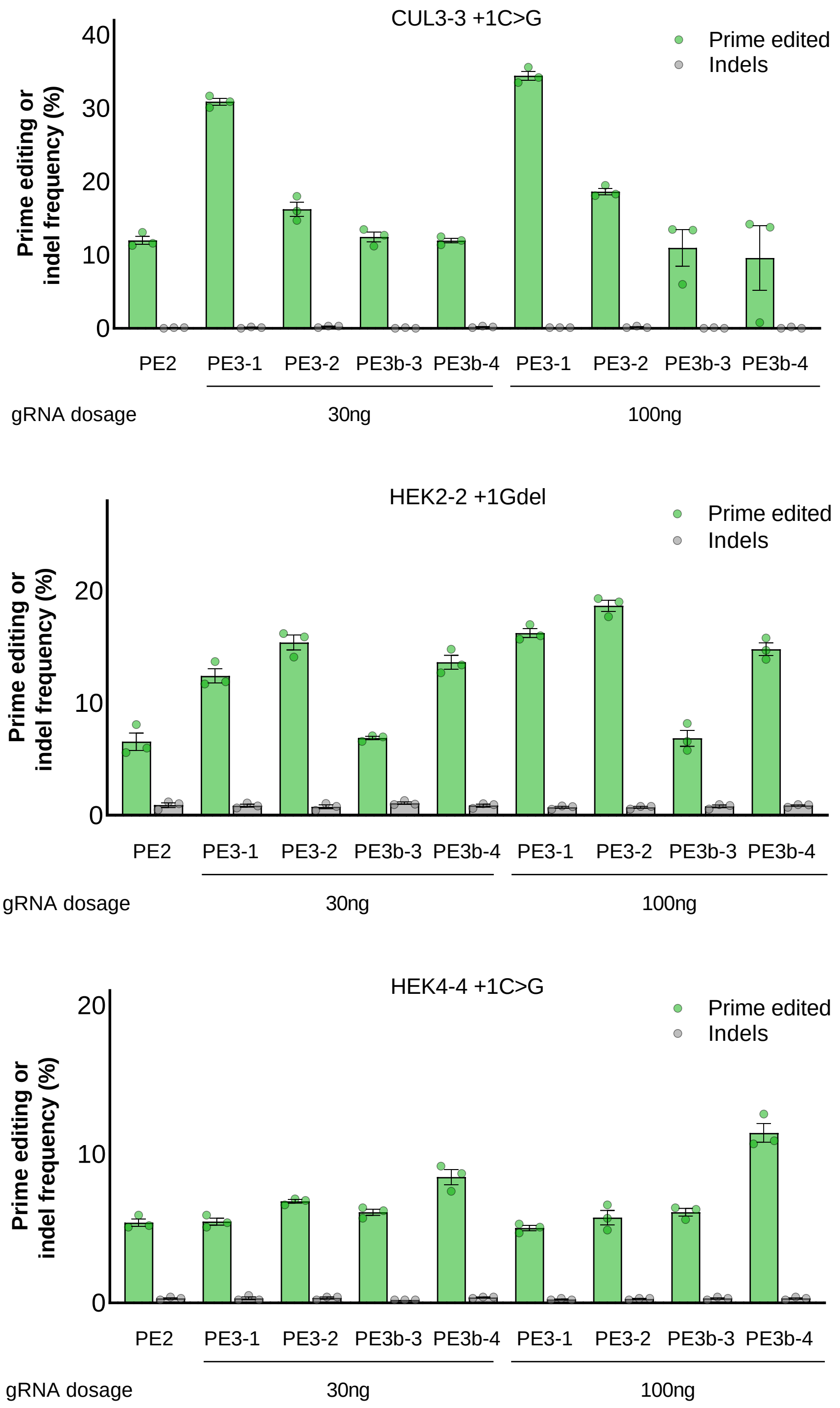


**Supplementary Figure 6.** Prime editing using PE3 and PE3b systems at three target sites. Two doses of gRNA were tested for PE3 or PE3b as described in the Method section and the prime editing activities with 30 ng gRNAs were plotted also in Figure 2a and 2b. There was no significant increase in prime editing activity or indels with the dose of gRNA across three target sites. The numerical values of the prime editing activity are described in Supplementary Table 1. Mean ± s.e.m. of $n = 3$ independent biological replicates.

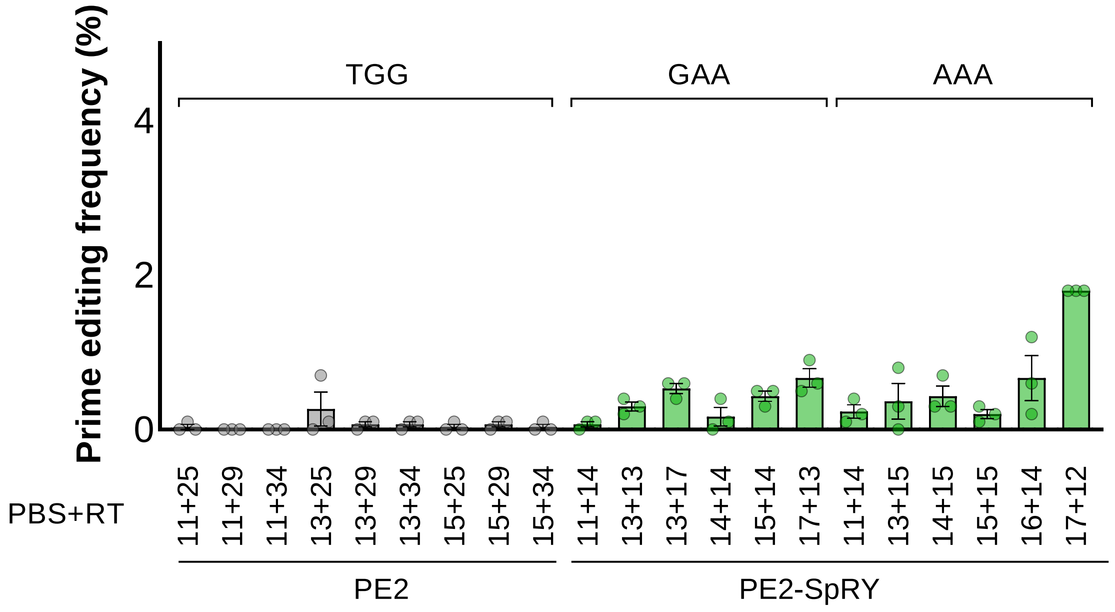


**Supplementary Figure 7.** Prime editing activities at the BRAF V600E site with pegRNAs of various lengths of PBS and RT template. The pegRNA of 13 nt PBS and 29 nt RT template showed the highest prime editing activity (average 0.3%) with wild-type PE2 and PE2-SpRY had average 1.8% prime editing activity with the pegRNA of 17 nt PBS and 12 nt RT template. The numerical values of the prime editing activity are described in Supplementary Table 1. Mean ± s.e.m. of $n$ = 3 independent biological replicates.

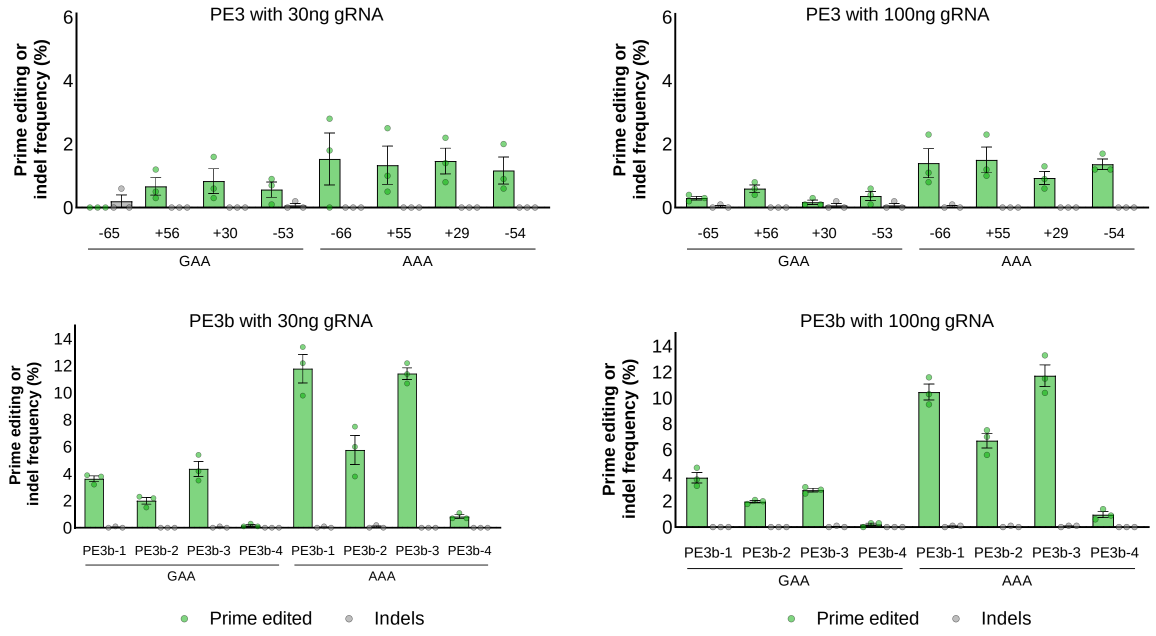


**Supplementary Figure 8.** Prime editing using PE3 and PE3b systems at the BRAF V600E site. The BRAF-GAA and BRAF-AAA pegRNAs were tested with four gRNAs in the PE3 system. Each gRNAs could induce nicks to the non-edited strand, and the nicked positions were indicated against the locations of pegRNA-induced nicks. The numerical values of the prime editing activity are described in Supplementary Table 1. Mean ± s.e.m. of $n = 3$ independent biological replicates.

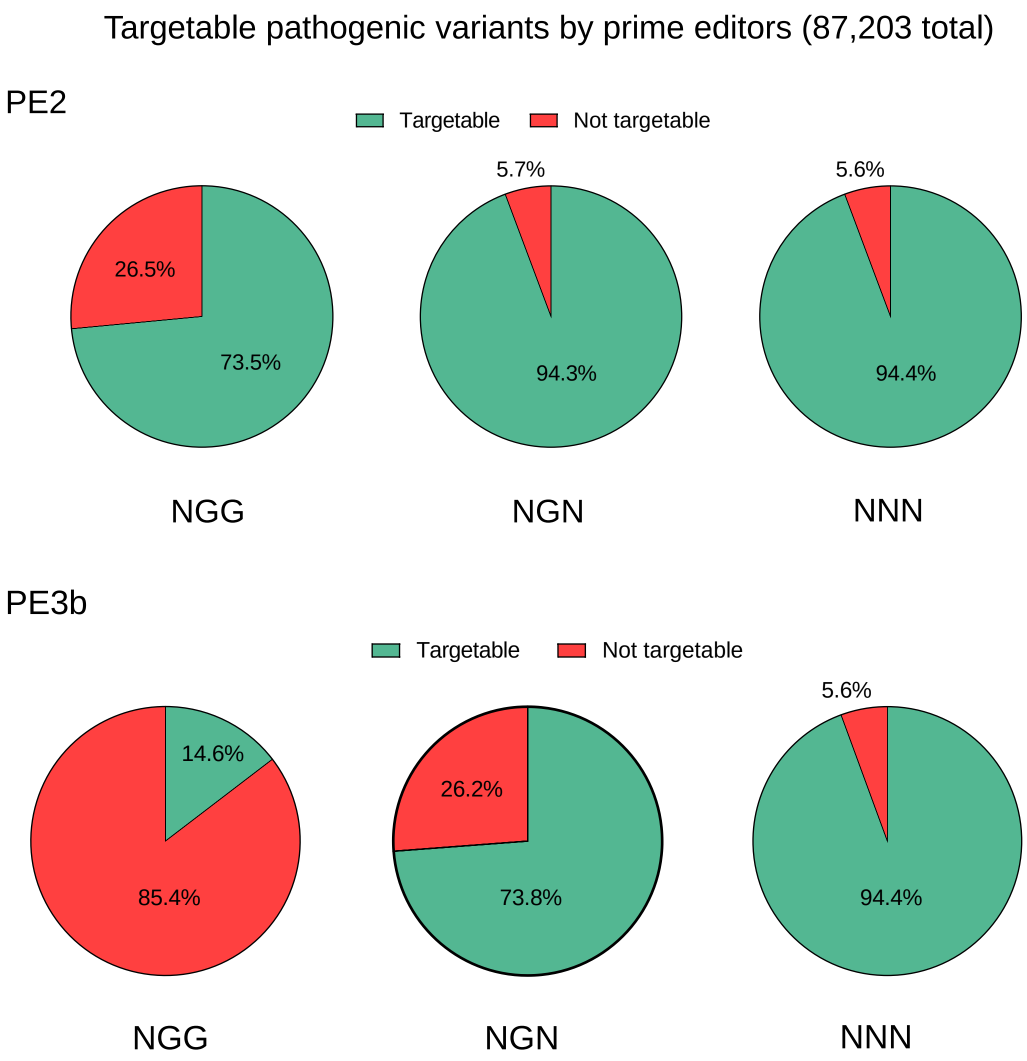


**Supplementary Figure 9**. Fraction of human pathogenic variants which could be targetable with the PE2 or PE3b system. 73.5% of pathogenic variants are targetable for wild-type PE2 (with NGG PAM) and 94.4% of pathogenic variants are targetable for PE2-SpRY (with NNN PAM). In the case of PE3b system, only 14.6% of variants are targetable by wild-type PE2, while the PE2-SpRY variant dramatically increases up to 94.4% of targetable pathogenic variants. The remaining 5.6% of pathogenic variants that cannot be targeted are large deletions, insertions, or complex mutations. Total counts are listed in Supplementary Table 4.

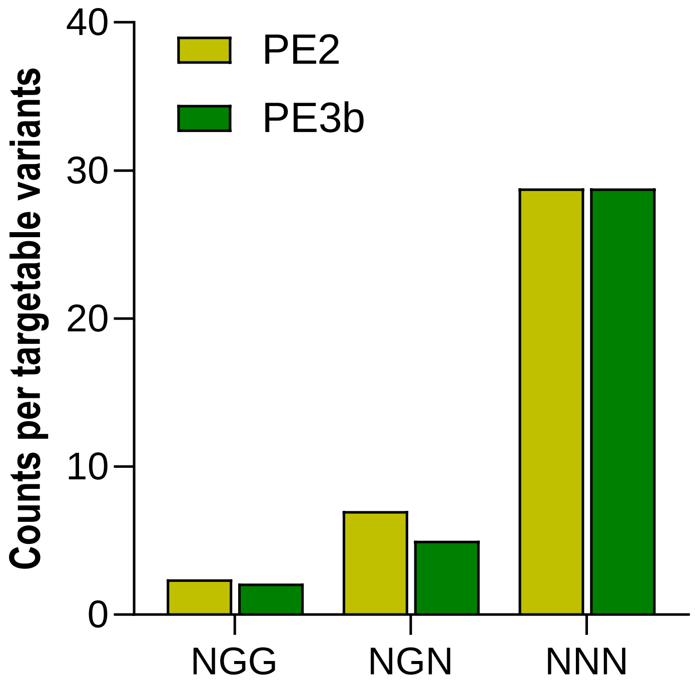


**Supplementary Figure 10**. Numbers of target site per targetable variants in the PE2 and PE3b systems. The PE2-SpRY variant increases the number of designable pegRNAs per pathogenic variant; from 2.4 to 28.8 pegRNAs in PE2 system and from 1.6 to 28.8 pegRNAs in PE3b system. Since the activity of prime editing is very diverse, the more pegRNAs that can be designed, the higher the success rate of prime editing. Total counts are listed in Supplmentary Table 4.

### Supplementary Table 3. List of off-target sites in this study.

| pegRNA names | | Chr. | Spacer sequences | PAM | PBS | RT template | Reference for off-target sites |
|---|---|---|---|---|---|---|---|
| HEK4 +2G>T | ON | Chr.20 | GGCACTGCGGCTGGAGGTGG | GGG | GCGGCTGGAGG | TTGGGGTTAA | Anzalone *et al*. |
| | OFF | Chr.10 | GGCACgaCGGCTGGAGGTGG | GGG | aCGGCTGGAGG | TTGGGGgTtg | |
| FANCF-4 +5Cins | ON | Chr.11 | GCAGAAGGGATTCCATGAGG | TGC | AAGGGATTCCATG | AGGTCGCGCGAAGGC | Walton *et al*. |
| | OFF | Chr.11 | GaAGAAGGGtTTCCATGAGG | AGA | AAGGGtTTCCATG | AGGaCGatacctgag | |
| EMX1-4 +5G>C | ON | Chr.2 | GTCACCTCCAATGACTAGGG | TGG | CCTCCAATGACTA | GGGTCGGCAACCA | |
| | OFF | Chr.17 | GTCACCTgtAATGACTAGGG | AGA | CCTgtAATGACTA | GGGaCaGtAAtgg | |
| MECP2-3 +1G>C | ON | Chr.X | GGGTGGTTCCATAATCTGTG | TAT | GGTTCCATAATCT | CTGTATACCTAAG | |
| | OFF | Chr.4 | GcGTGGTTaCATAATCTGTG | GAG | GGTTaCATAATCT | CTGgAggggTgca | |
| CUL3-3 +1C>G | ON | Chr.2 | GAGTTCTCATGGAGTGACTG | CTC | TCTCATGGAGTGA | GTGCTCACGTAAC | In this study |
| | OFF1 | Chr.14 | aAGTTtTCATGGAGTGACTG | TCA | TtTCATGGAGTGA | GTGtcacaGagta | |
| | OFF2 | Chr.2 | GtGTTaTCATGGAGTGACTG | AGA | TaTCATGGAGTGA | GTGagaAtGacAC | |
| | OFF3 | Chr.8 | GAGTTCTgATGGAGTGACTG | CTG | TCTgATGGAGTGA | GTGCTggtGccca | |
| HEK2-4 +1Cins | ON | Chr.5 | GGGCGGGCCAGCCTGAATAG | CTG | GGGCCAGCCTGAA | CTAGCTGCAAACAA | |
| | OFF1 | Chr.13 | GGGCtGGCCAGCCTGAtTAG | AAT | tGGCCAGCCTGAt | CTAGaataggctAA | |
| | OFF2 | Chr.22 | GGGCGaGCCAGCCTG(g)AATAG | GTA | GaGCCAGCCTG(g)AA | CTAGgTaCtggCct | |
| | OFF3 | Chr.5 | aGGCtGGCCAGCCTGAATAG | CTT | tGGCCAGCCTGAA | CTAGCTtCAgcagg | |

The mismatches are highlighted in blue and lower case. At the HEK2-3 +1Cins OFF-2 site, () means a bulge position compared to the on-target site. The intended mutations of pegRNAs are shown in red.

**Supplementary Table 4.** Analysis of targetable pathogenic variants using PE variants.

| Types | | Number of targetable variants | | number of pegRNAs per target variants | |
|---|---|---|---|---|---|
| **Prime editors** | **PAM** | **Counts** | **Percentage (%)** | **Total counts** | **pegRNAs per targetabel variants** |
| **PE2** | **NGG** | 64,121 | 44.8% | 156,862 | 2.4 |
| | **NGN** | 82,239 | 79.2% | 572,093 | 7.0 |
| | **NNN** | 82,283 | 89.6% | 2,372,052 | 28.8 |
| **PE3b** | **NGG** | 12,769 | 7.9% | 26,589 | 2.1 |
| | **NGN** | 64,315 | 51.3% | 320,763 | 5.0 |
| | **NNN** | 82,283 | 83.8% | 2,372,052 | 28.8 |

*Total of 87,203 variants from ClinVar database were analyzed.
